# MAGELLAN: a tool to explore small fitness landscapes

**DOI:** 10.1101/031583

**Authors:** S Brouillet, H Annoni, L Ferretti, G Achaz

## Abstract

In a fitness landscape, fitness values are associated to all genotypes corresponding to several, potentially all, combinations of a set of mutations. In the last decade, many small experimental fitness landscapes have been partially or completely resolved, and more will likely follow. MAGELLAN is a web-based graphical software to explore small fitness/energy landscapes through dynamic visualization and quantitative measures. It can be used to explore input custom landscapes, previously published experimental landscapes or randomly generated model landscapes.

## 2 INTRODUCTION

Sewall Wright (1932) first introduced fitness landscapes as a metaphor to study evolution. Fitness landscapes have been increasingly popular in the last couple of decades (recent reviews by Orr (2005) and de Visser and Krug (2014)) as more and more landscapes were experimentally resolved (see Table 1 in Weinreich et al., 2013). Fitness landscapes were not only a cornerstone in our understanding of evolution (Maynard Smith, 1970; Kauffman, 1993; Gavrilets, 2004) but also contributed to the scientific exchange with other fields, especially with computer science (Richter, 2014) and physics (Stein, 1992).

A fitness landscape is a complex multidimensional object that maps each genotype, *i.e.* a combination of alleles hosted at different loci, to a fitness value. In experimental fitness landscapes, the fitness is assessed through a proxy (growth rate, antibiotic resistance, etc.) that is supposedly proportional to the fitness. In the energy landscapes described in physics literature, the fitness values are replaced by energy values but the overall object is identical. For the rest of this article, we will only use population genetics terminology (loci, genotypes, fitness, etc.) for the sake of clarity, but we would like to emphasize that MAGELLAN can be equally used to explore any type of landscapes: model fitness landscapes with properly defined fitness, experimental fitness landscapes with proxies or energy landscapes from physics.

The genotypes are composed of several polymorphic loci with two or more alleles. When restricted to bi-allelic loci, the genotype space is a hyper-cube of size *2^L^,* with *L* the number of loci. More generally, the genotype space is discrete and has a size of 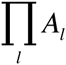, where *A_l_* is the number of alleles at locus *l*. Each genotype is then associated to a fitness value, typically a real number representing the relative reproductive success.

Because the size of fitness landscapes grows exponentially with the number of loci, their visualization and systematic exploration is essentially impossible for landscapes of more than several loci. In practice, only complete landscapes of typically 10–15 loci can be reasonably analyzed on a modern computer: simply storing fitness values of a bi-allelic landscape of 15 loci requires 1Gb of memory (using floats of 4 bytes). Although these landscapes seem small when compared to a genome-size fitness landscape (millions or even billions of 4 allele loci), the analysis of small fitness landscapes remains crucial to study evolutionary processes in a small set of polymorphic loci with genetic interactions.

It is noteworthy to mention that other fitness landscapes based on phenotypes were also defined. Phenotype-based fitness landscapes, such as the popular Fisher geometrical model (Fisher, 1930; Tenaillon, 2014), associate phenotypes (instead of genotypes) to fitness values. Phenotypes are usually characterized by *T* independent continuous real traits, resulting in a phenotype space of dimension ***R****^T^.* However, as the current implementation of MAGELLAN only handle genotype-based fitness landscapes, we will omit phenotype-based fitness landscapes for the rest of this article.

Ideally, one would like to analyze the structure of the fitness landscapes independently of any evolutionary processes. However, even the definition of a fitness landscape has hidden assumptions on the evolutionary processes. For example, having a fixed landscape regardless of the frequencies of the genotypes assumes that the fitness is frequency-independent, a very strong but often useful assumption. Classical quantitative measures of landscapes (*i.e.* summary statistics) are also often defined to characterize some evolutionary processes. For example, the length of fitness increasing paths is meaningful only in a model where the population is genetically homogeneous and is abstracted as a single particle that climbs the peaks of the landscape (the so-called “strong selection weak mutation” regime defined by Gillespie (1983)). In experimental fitness landscapes, the fitness proxies can only be considered as good substitutes for genuine fitness under some strong assumptions and even the scale (linear, log or exponential) where the landscape should be analyzed could be hard to define.

In the original Wright representation, all genotypes are located on a flat plan and a third dimension is used for fitness (Wright, 1932). These metaphoric fitness landscapes are aesthetically appealing (see e.g. Figure 2 in Orr, 2005), but cannot be used to properly study fitness landscapes. Indeed, because of the high dimensionality of the neighborhood, genotypes cannot be placed on a flat plane while keeping the correct distances between them. This simple argument gave rise to criticisms against the usefulness of fitness landscapes visualization (e.g. Provine 1986, Gavrilets 2004). Quite correctly, only a two-locus two-allele landscape can be properly mapped on a plan.

Therefore, representing visually fitness landscapes with more than 2 loci is a challenging problem. At least two ideas were previously suggested. Wiles and Tonkes (2006) took advantage of a recursion to systemically split the hypercube into squares that are connected by their corner positions. They then represented the whole hypercube on a square matrix. Although, this representation has explicit connections between the neighboring genotypes, it requires some training for navigation. It is furthermore difficult to assess the properties of the landscape in this representation. For example, testing visually for the additivity of fitness is almost impossible. McCandlish (2011) proposed another visualization that is explicitly based on the evolutionary processes. The author used the main axes of a PCA-like decomposition of the transition matrix between genotypes to place genotypes on a plan. As the transition rates depend non-linearly on the population size, the proximity of genotypes and therefore the overall representation depends on the population parameters, especially population size. This last representation is well suited to explore large fitness landscapes. We however believe that an alternative simpler representation would be helpful in deciphering the structure of small landscapes.

MAGELLAN aims at giving a visual representation of a small fitness landscapes (up to 10 loci) and at providing tools to analyze them. Indeed, as the visual representation is doomed to be approximate, characterizing the structure of a fitness landscape is a major challenge. It often relies on summary statistics that quantify *a priori* chosen properties of the landscape. MAGELLAN generates several complementary views of a given landscape and computes systematically the set of summary statistics that were proposed in the literature (reviewed in Szendro et al., 2013; Ferretti et al., submitted). We believe that the joint use of visual representations together with the analysis of summary statistics is the key to unravel the structure of small landscapes.

## 3 DESCRIPTION OF MAGELLAN

MAGELLAN (MAps of GEneticaL LANdscapes) is an intuitive and simple visual web-based representation especially designed to explore small genotype-based fitness landscapes, typically less than 10 loci. It can be used to explore model and experimental fitness landscapes. More precisely, it generates through simulations and analyzes model fitness landscapes; it browses published experimental landscapes or it takes an input custom landscape. For a given landscape, MAGELLAN will create a visual representation that can be dynamically rotated or tuned by several options, and compute the summary statistics that characterizes it.

### 3.1 Visual representation

In the standard view (Figure 1), genotypes are sketched as sequences of circles filled with colors that represent the alleles. A genotype’s position is set on the x-axis by its Hamming distance to a reference genotype (placed at x=0) and on the y-axis and by its associated fitness. A genotype is connected to its neighbors by green, red or orange lines that represent fitness gains, losses or neutral mutations from the wild-type allele. The background of peaks (genotypes with no fitter neighbor) is highlighted in green, and in red for sinks (genotypes with only fitter neighbor).

**Figure 1.**
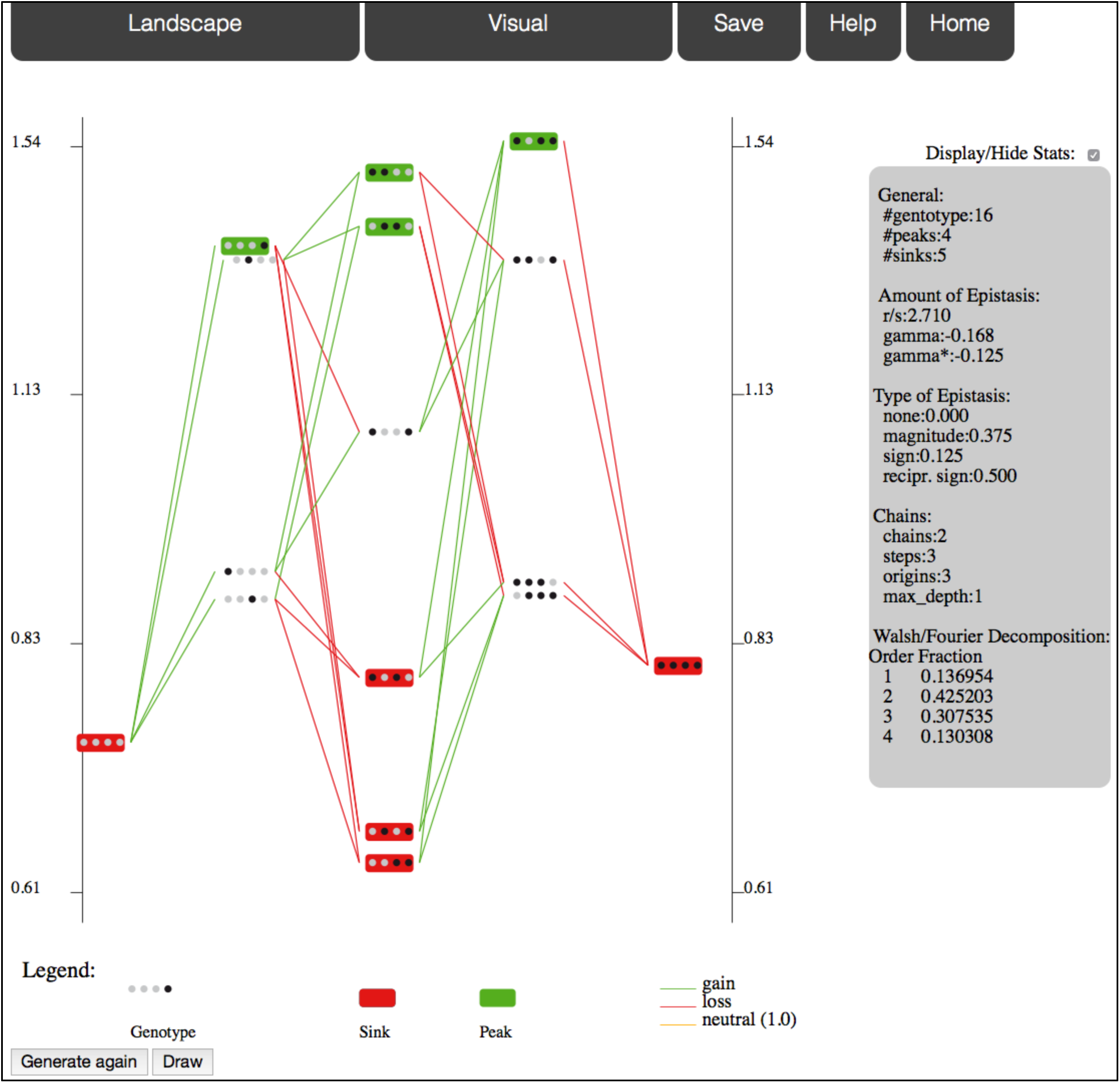
One standard view of a House of Cards model fitness landscape with 4 loci of 2 alleles.

The main pages contain menus that are arranged in several tabs, among which ***Landscape*** to specify the current landscape and ***Visual*** to set options on the visual representation. By default, all summary statistics are displayed on a right of the representation. Once a representation is displayed, users can tuned on and off several options and then update the display by clicking ***Draw***. All options are accessible under the ***Visual*** tab, but some can be directly changed from the graph. For example, simply clicking on a genotype chooses it as the new reference genotype. Other options include log-scale, path selection (starting from and/or ending at a genotype), zoom, threshold ratio for neutrality, mutation at a single locus, sub-landscape, etc.

Users can also change the view to ***Compact***, shrinking genotypes to small squares (losing the allelic sequence) and/or to ***Flat***, to shows a view from the sky, that is closer to Wright original drawing. We think that the flat view is especially interesting when only a subset of the paths is displayed (e.g. increasing fitness paths from/to a genotype, chain trees only or large jumps in fitness). Alternating between the different views and rotating the landscape is essential to get a visual exploration of the landscape and to appreciate the meaning of the summary statistics.

Incomplete landscapes are displayed without the missing genotypes.

### 3.2 Summary statistics

MAGELLAN reports, on the panel located at the right of the representation, the complete set of summary statistics that were proposed in the literature:

- Number of **peaks** (Weinberger, 1991), which are genotypes with no fitter neighbor. It also corresponds to the number of local fitness maxima.
- Number of **sinks** (Ferretti et al., submitted), which are genotypes with only fitter neighbors.
- **r/s ratio**, --roughness/slope-- (Aita et al., 2001): the landscape is fitted to a linear model (a linear combination of independent fitness effects plus a constant) by least squares. The slope s is the average fitness effect, whereas the roughness *r* is the quadratic mean of the residuals of the regression. The larger this ratio, the more noise (*i.e.* epistasis) in the landscape.
- **Types of epistasis** (Weinreich et al., 2005; Poelwijk et al., 2007): fraction of pairs of loci that have no epistasis, magnitude epistasis (change in fitness effect without change in sign), sign epistasis (one of the two mutations has an opposite effect in both backgrounds) and reciprocal sign epistasis (both mutations have an opposite effects on the other background). Note that the definition of sign epistasis and reciprocal sign epistasis does not depend on the scale (e.g. linear or log) whereas the magnitude epistasis does.
- Amount of epistasis assuming a **Fourier expansion** of the landscape (Stadler, 1996; Weinreich et al., 2013; Neidhart et al., 2013). It is the fraction of interactions that cannot be reduced to simple additive fitness. We report the fraction of interactions of order 2 as well as the interactions of higher order.
- γ and γ^*^ (Ferretti et al., submitted): correlation in fitness effects between genotypes that only differ by 1 locus, averaged across the landscape. γ^*^ is the correlation in sign and is therefore independent of the scale (linear or log).
- Number of **chain** trees, chain steps and chain depth (Ferretti et al., submitted): chain steps are the genotypes with a single fitter neighbor and chain depth gives information on their relative arrangement.

### 3.3 Model landscapes

Several classical models of fitness landscapes are implemented. They are first specified by a number of loci and alleles. Random values are then drawn from normal or uniform distributions, using the input parameters. The ***Generate again*** action redraws all random values and displays a new realization of the model, while keeping all selected options identical. Generating several landscapes with identical parameters gives a glimpse at the diversity of landscapes that can be generated. The models currently implemented are:

- **Multiplicative**: All non-wild-type alleles have a 1+s independent fitness contribution, where s is a normal random variable. This model has no epistasis and is also known as the additive model (in log-scale, products become sums).
- **House of Cards**: (Kingman, 1978) the log-fitness of all genotypes are i.i.d. random variables from a normal distribution with mean 0.
- **RMF**: (Aita et al., 2000) Rough Mount Fuji is the sum of a House of Cards with a Multiplicative model. The relative contribution of both is tuned by the relative values set for the Multiplicative and the House-of-cards parts.
- **Kauffman NK**: (Kauffman and Weinberger, 1989) Each locus interacts with K other loci and contributes by a uniform [0,1] random fitness that depends on the state of all its interacting loci.
- **Ising**: (Mézard et al., 1987) Each locus interacts with both its left and right neighbors. For each pair of interacting loci, there is an associated random log-fitness cost if the alleles at the two loci are different.
- **Eggbox**: Each genotype has either high or low random fitness, with a large jump in fitness between neighbors.
- **Full Model**: a linear combination of the above models.

### 3.4 Implementation

MAGELLAN was written in standard C. It is based on an open access library of functions that generate and analyze fitness landscapes. A command line version of the program as well as the library is available upon request. The web-based implementation uses HTML5 specifications and java-scripts (that are handled in all recent browsers). The authors will keep updating MAGELLAN and incorporate any interesting new features suggested by the users.

## 4 CONCLUSION

MAGELLAN provides an easy-to-use graphical interface to explore both experimental and model fitness/energy landscapes. As these landscapes are highly dimensional, there is no single 2-dimensional representation that can capture well their structure. Therefore, we encourage the users to confront the analysis of summary statistics to several visual representations, obtained by changing the reference genotype or by trying the flat and compact view. MAGELLAN will keep updating its experimental and model landscapes, its summary statistics and will incorporate interesting suggestions. One direction that we are willing to pursue is the superposition and/or comparison of multiple landscapes on the same genotype space.

## 5 ACKNOWLEDGEMENTS

The authors would like to thank O. Tenaillon for early input on the display of landscapes, J. Krug for suggestions of new features on a previous version of the program and D. Weinreich for his feedbacks and scientific input. This work was supported by the ANR grant TempoMut ANR-12-JSV7–0007.

